# PuMA: PubMed Gene-Celltype-Relation Atlas

**DOI:** 10.1101/2024.02.09.579584

**Authors:** Lucas Bickmann, Sarah Sandmann, Carolin Walter, Julian Varghese

## Abstract

Rapid extraction and visualization of cell-specific gene expression is important for automatic celltype annotation, e.g. in single cell analysis. There is an emerging field in which tools such as curated databases or Machine Learning methods are used to support celltype annotation. However, complementing approaches to efficiently incorporate latest knowledge of free-text articles from literature databases, such as PubMed are understudied. This work introduces the PubMed Gene/Celltype-Relation Atlas (PuMA) which provides a local, easy-to-use web-interface to facilitate automatic celltype annotation. It utilizes pretrained large language models in order to extract gene and celltype concepts from PubMed and links biomedical ontologies to suggest gene to celltype relations. It includes a search tool for genes and cells, additionally providing an interactive graph visualization for exploring cross-relations. Each result is fully traceable by linking the relevant PubMed articles. The software framework is freely available and enables regular article imports for incremental knowledge updates. GitLab: imigitlab.uni-muenster.de/published/PuMA

## 1 Introduction

In the field of single-cell RNA-sequencing (scRNA-seq) analysis, the interpretation of the large variety of gene markers, celltypes and their specific (co)-expression profiles have been a challenge for many years. Manual annotation is time-consuming and partially subjective (1). However, as of today, manually curated marker-lists are still used to identify and annotate cells in scRNA-seq analysis. Attempts are made to tackle this approach by databases, such as *CellMarker2* (2) and *PanglaoDB* (3). CellMarker2 is curated manually and aims at covering a range of markers based on high quality published articles. Still, it relies on structured input, and is not automatically updated based on newest research literature. CellMarker2, therefore is limited by covering only 24591 studies. PanglaoDB, analog to CellMarker2, is a database for exploring and analyzing scRNA-seq data from mouse and human tissues. It contains pre-processed and precomputed analyses from more than 1054 single-cell experiments, with more than 4 million cells from a wide range of tissues and organs. The authors have also compiled a manually curated list of more than 6000 markergenes, which can be used for celltype annotation. However, the database is no longer supported, and the collection of marker-genes is manually curated as well.

This outlines the importance and potential of Biomedical Named Entity Recognition for research and discovery, which is facilitated through machine learning in the recent years (4). Advancements in the field of natural language processing (NLP) emerged successfully with new deep learning techniques such as large language models (5) [Quelle]. Tools, such as *MarkerGenie* (6) and *Advanced Biomedical Entity Recognition and Normalization 2* (BERN2) (7) have been fine-tuned to the domain of biomedical texts. This facilitates the annotation of biomedical entities for large amounts of articles. In this work, we present PuMA - a regularly updated gene/celltype-relation database based on PubMed. A locally deployable Docker application provides an easy-to-use interface and an interactive graphical visualization of gene/celltype-relations.

## 2 Methods

PuMA is created, updated, and provided with two independent Docker containers. 1) the Updater, which is necessary to integrate new articles from PubMed, 2) a Webserver, used for querying and viewing the results. We also provide the user with easy access of regular updates, based on latest published articles. We therefore have a two-fold approach: We provide a downloadable updated database releases in regular schedules. We also provide the Docker container for updating the database, running independently from our provided infrastructure. In the following, the database creation and querying are explained in detail.

### 2.1 Atlas creation scheme

A complete workflow is presented in Figure 1. The annotations of PubMed articles are extracted by BERN2, a natural language model fine-tuned for biomedical named-entity recognition applications. It is capable of annotating celltypes, genes, cell-lines, species, mutations, drugs, and diseases. We incorporate this model to annotate newly published articles, regarding celltypes and genes. We process and filter all annotations returned by BERN2, their classification as ‘celltype’ or ‘gene’ and their mapped normalized Identifiers (IDs) from the *Cell Ontology (CL)* (9) and *NCBI Gene* (8) databases.

**Figure 1.**
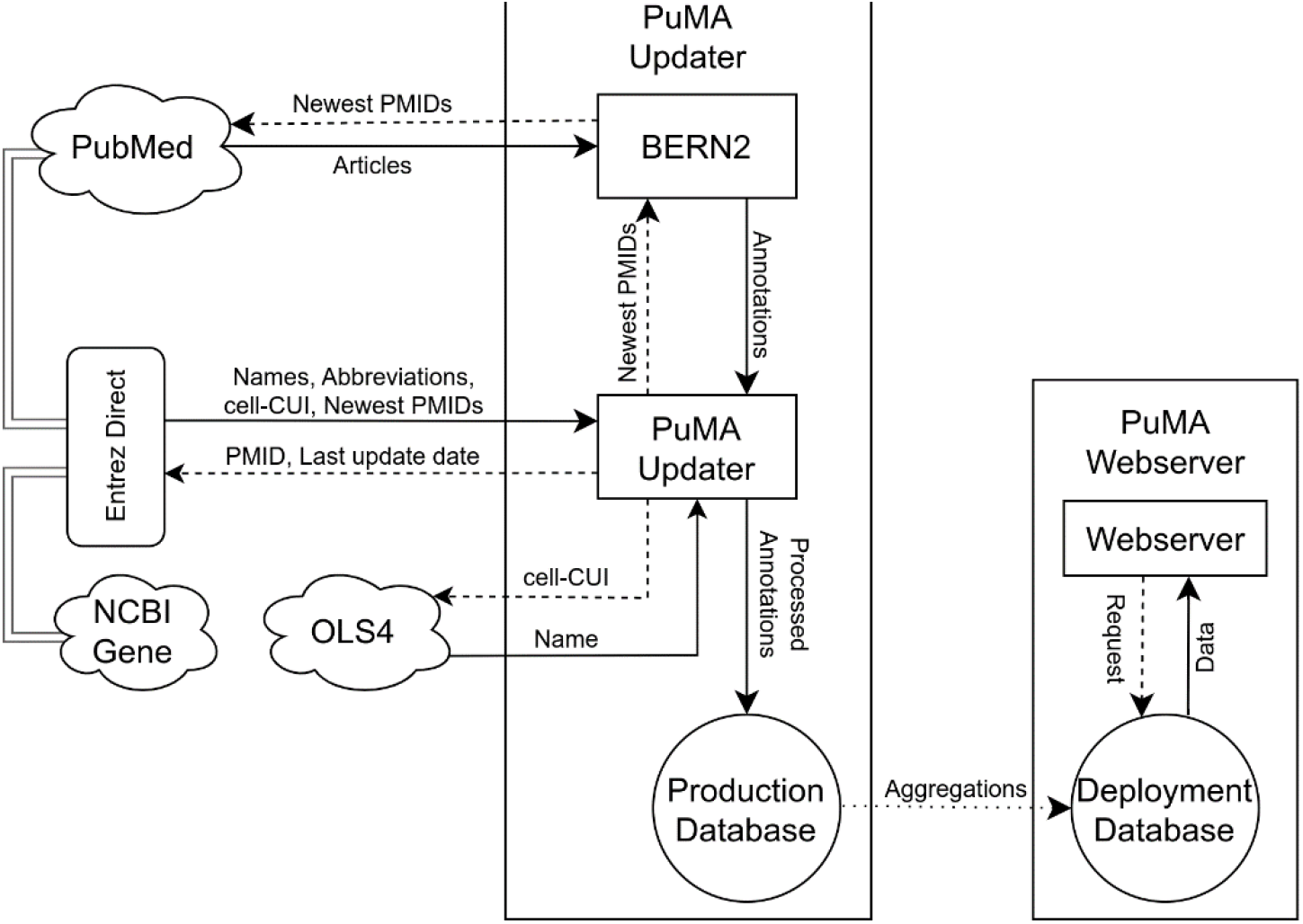
PuMA creation schema. PuMA consists of an Updater (middle) and the deployable Webserver (right).

The *Entrez Direct* interface by the National Center for Biotechnology Information’s (NCBI) (10) is used for retrieving meta-information, such as journal-names, publication-dates and the PMIDs of newly published articles. Additionally, the concepts unique identifiers (CUIs) of genes are looked up in NCBI Gene. The CUIs are maintained by the Unified Medical Language System (11) and helpful to identify common biomedical concepts or generate semantic core datasets (12). The European Molecular Biology Laboratory’s European Bioinformatics Institute’s *Ontology Lookup Service (OLS) version 4* (13) is used to retrieve standardized names for the CUI of annotated cells, to reduce the highly variable spellings returned by BERN2. All information is stored in a production database. After successful storing of new annotations, the database is pre-filtered and aggregated into a deployment database. This process involves the removal of relations based on low evidence and co-occurrences. The computation is based on the number of articles referencing the same gene to celltype relation. Relations with low co-occurrences per article or few overall referencing articles are filtered. This reduces its deployment size, while maintaining all relevant information to the user. The updated deployment database is provided regularly, and can be downloaded independently aside the local running webserver (see section 2.2), to allow independent updates. The Updater Container is only required if a user wants to update and maintain the deployment database him-/herself.

#### Detailed Workflow

The production database keeps track of the last date, at which released articles were processed. PubMed is queried via Entrez Direct to retrieve all newest PMIDs and articles. Subsequently, these are processed by multiple local dockerized BERN2 instances. The output, annotations for each new article, is processed in a concurrent manner. An additional filtering step, to remove annotations without mapped gene-or cell-CUI, is conducted alongside. Additional information, such as the mapping of the CUI to official names and their abbreviations are retrieved via Entrez Direct for genes and OLS4 for cells. Optionally, information on the journal and the publication date can also be retrieved via Entrez. Redundant information, such as duplicate cell-names for multiple occurrences in several PMIDs, is removed by using a normalized database form. An updated version of the production database, containing all newly added annotations, is stored.

A deployment database is created by aggregating information from the production database by grouping same cells and genes by their CUI. In PuMA, several statistics are calculated, which include, e.g. the number of distinct PMIDs (*pmid_counts*), and the derived total occurrence count of a gene/celltype-relation (*total_counts*). Additional filtering of few co-occurrences and general occurrences (≤ 3 PMIDs), and SQLite optimizations results in a smaller, more efficient database for the user, while retaining all information in the global production database. Additional updates are therefore still incremental and depend on the number of new annotations. The user updates to a new release with a simple download of the updated deployment-database from the docker registry.

### 2.2 Webserver, querying and usability aspects

The webserver is a standalone Docker container, which provides all necessary interfaces on a web application to query the provided database. This includes, among others, a search field, filter for exact matches, and a cutoff-parameter for low-scoring hits. Any user query is locally processed. Based on the requests on the frontend (such as a gene- or cell-search), the query-term and search-parameters are sent to the backend, after which a specific SQL-query is executed on the deployment-database. Hits are processed, including the aggregation of found PMIDs and the computation of the scores for each entry, before being sent to the frontend for visualization. These scores are calculated for each gene/celltype-relation using a variation of log normalized frequency-inverse-document-frequency (TF-IDF) (see Equation 1). The logarithm reduces the scale of extremely large numbers, while multiplication ensures that the number of supporting articles is weighted by the total number of occurrences. Logarithmic scoring is more robust against outliers than a linear function, and the computation is easy and efficiently implementable for the SQLite backend. In contrast, the commonly used TF-IDF (14), with inverse-document-frequency, would filter out common genes and celltypes, such as B-or T-cells, and would be prone to boost lower occurring subtypes. This is because IDF gives more weight to rare terms, which leads to the deranking of often studied and commonly occurring celltypes, which is undesired. Instead of using IDF, we suggest scaling the logarithmic frequency of the document frequency itself. This approach preserves the original distribution of the data and ensures that all terms are treated equally. We apply this transform to score hits, and not to extract these by studying its occurrences.

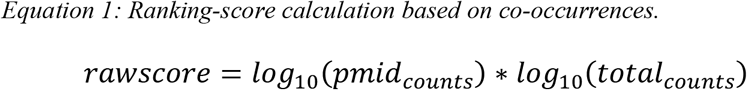

Subsequently, the scores are normalized, by dividing all scores by the top-3 hit. This improves usability aspects of the application, by reducing information overflow and generate an independent unit of importance measurement. Therefore gene/celltype-relations with higher normalized scores are clearly identifiable and comparable for the same query, and all values now apply to the relative importance to each other. As the dataset is large, it usually has a high-importance outlier and a larger number of low-scoring hits. This scaling ensures that low-scoring hits can be easily cutoff, and the top-1 hit does not distort the score distribution of the average scoring results. These normalized tables are presented to the user on the web-interface. Additional functions have been implemented, e.g. searching, sorting, exporting the table. All relevant PubMed-articles for each result are listed and linked in a separate interface, which allows full transparency and explainability of the results. The listed PubMedIDs are buttons, linked to their corresponding PubMed entries, and open the respective article in a new tab. This leads to a highly explainable tool, where all relevant articles are linked for the user, and which can be accessed for each hit individually. Table 1 shows an exemplary result for a gene search for the search term ‘A1BG’ sorted by descending score.

**Table 1:**
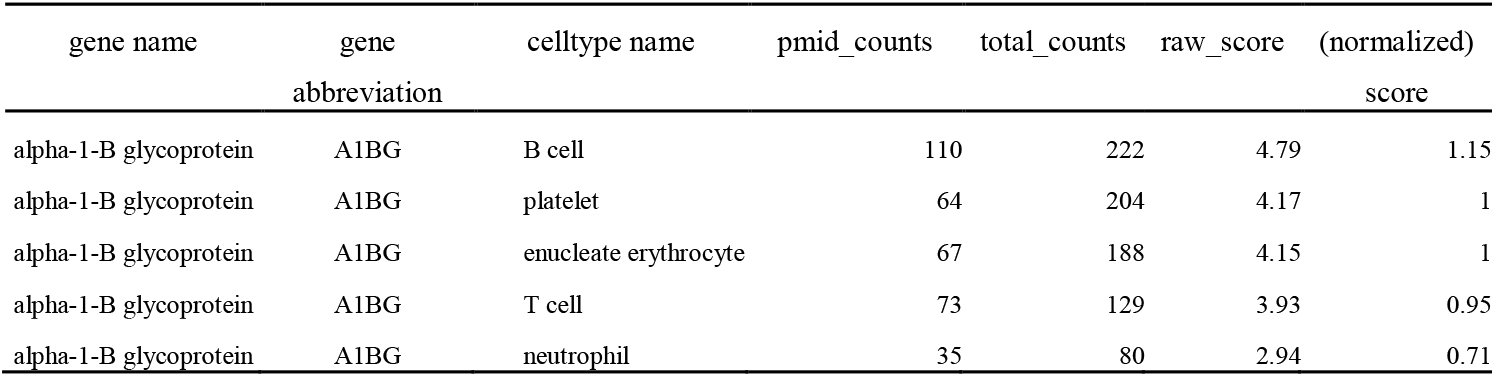
Extended database result for search-term ‘A1BG’. The table is sorted by descending (normalized) score.

#### Graph construction

A graph is computed based on the same workflow as previously described. The only difference is that the search-term that is applied for genes and cells concurrently, and the results are merged. A graph is constructed by representing celltypes and genes as nodes, and relations as corresponding edges, based on the gene/celltype-cooccurrence in PubMed articles. The node-sizes and edge-widths are both scaled by their normalized score, using their highest corresponding entry. The *Compound Spring Embedder Layout (CoSE)* (15), a force-directed graph layout, is used to present the graphs. Gene-nodes are presented with their abbreviation for clearer visibility. Their full name can be shown by selecting the corresponding node. An example for a search-term ‘corn’ can be seen in Figure 2. The figure presents two celltypes and their corresponding co-occurring genes. They share three genes, which indicates a cross-regulation between these celltypes. The size of the node and edge indicate the supporting evidence, such as ‘epidermal growth factor’ has a higher correspondence on the ‘corneal epithelial cell’ than ‘interleukin 6’ (IL6), or the ‘corneal endothelial cell’. These cross-interactions between genes and different cells can be explored interactively using the graph visualization of PuMA.

**Figure 2.**
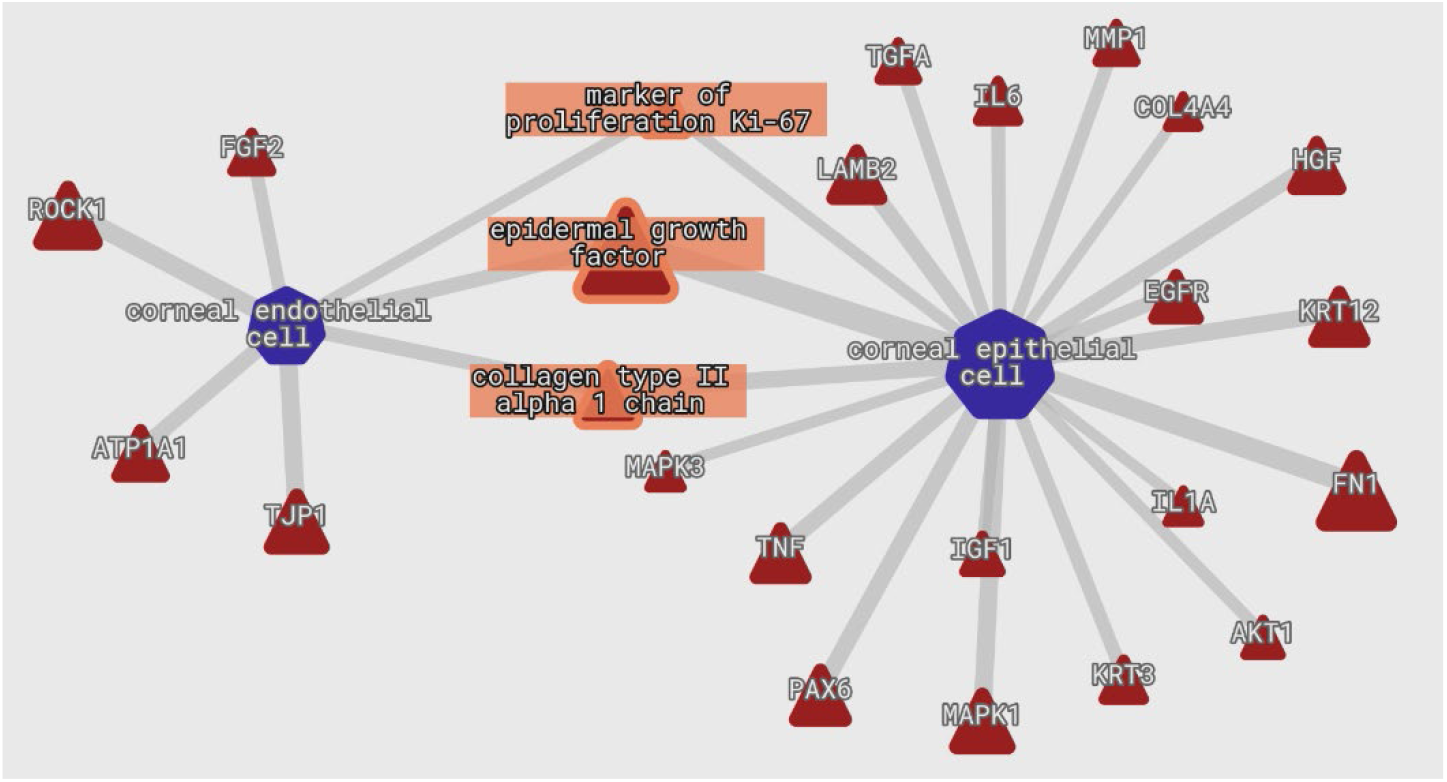
Cropped graph example for the search-term ‘corn’. Genes and celltypes are represented by red triangles and blue heptagons respectively. Nodes and edges are scaled by their normalized score. Overlapping genes have been selected and are therefore highlighted.

#### Application Interface

The tool also provides an API-endpoint for python projects. It allows for easy access of the same database using a python class and interface. The same functionality is given with exception of the graphical visualization and the combined query used for the graph search. This API can be installed via a python package manager, such as pip - using a local install inside the cloned PuMA repository.

### 2.3 Evaluation

We conducted an evaluation of query results based on two scRNA-seq datasets with known celltypes. First, a mouse dataset, *Tabula Muris (TM)* (16), and second, a human *Peripheral Blood Mononuclear Cells (PBMC)* (17) dataset. The evaluation is conducted in comparison to CellMarker2, a manually curated database, with experimentally supported markers of various celltypes. The scRNA-seq datasets were chosen for being well-characterized, publicly available and consider two relevant species – mouse and human. We compare the results generated with CellMarker2 to our work, PuMA. CellMarker2 is chosen, as it is one of the largest and latest curated databases available. We evaluate both approaches by computing of a unified distance score (see next section) for each of the top 50 differentially expressed genes (DEG), taken from Shunfu Mao et al. (18). The evaluation conducted by the authors involved using clusters of gene expression and reference celltype annotations obtained from the TM and PBMC papers. For each cluster, they extracted the top n = 50 DEGs by 1-versus-rest gene expression ratio. We chose all 50 genes per celltype, which are considered markergenes, as this number is specific enough for a broad analysis, and higher than manually analysis could provide in a reasonable time window. We analyze the computed distance scores with a top-1 hit per gene evaluation. Three statistics were evaluated: 1. Number of queried genes with existing gene-celltype relation, 2. Compared distance performance and 3. The combined performance. For the first, we searched each distinct DEG against both databases. Any celltype hit corresponds as a matched gene. A Venn-Diagram visualizes the occurrences of these in either, both or none of the databases. The compared distance is evaluated, by computing the distance of the top-1 matched gene/celltype-relation to the respective annotated reference celltype. The plot is created by visualizing the mean difference over all genes with a confidence-interval, as not all genes score equally well. The combined distance is the lower top-1 hit distance for either database for each gene respectively. It is visualized analog to the direct distance comparison. The scripts for evaluation are provided in the supplementary (see README for details).

We apply filtering to both databases (CellMarker2 and PuMA), to keep only information about gene/celltype-relations, which mention at least 3 articles each. We encourage to apply this filtering, as at least three individual resources should be the bare minimum as valid basis for profound information, even for manual curated databases. We applied species filtering for CellMarker2 to increase its sensitivity.

As CellMarker2 is a curated database, we expect it to perform better in certain cases. However, as its size is largely limited, the combination of curated and an extensive database, in this case PuMA, should increase the number of found genes and cells substantially. Therefore, we also analyze the capabilities of combining databases, and applying a top-1 evaluation with both databases combined. We query both for the same marker genes and use the combined minimum distance for each result. Given any number of results not present in both databases, we use its singularly corresponding top-1 hit. The statistics are given with a 1 sigma confidence interval (68%) if applicable.

#### Distance Evaluation

We evaluate the ‘correctness’ of annotated vs reference celltypes by the scRNA-seq datasets, using a standardized scoring system based on the Cell Ontology database. Based on the *v2023-07-20* release, we traverse its directed acyclic graph with a modified Dijkstra algorithm. Parent and child relations, such as ‘anucleate cell’ and ‘corneocyte’, increase the score by one, siblings, such as ‘corneocyte’ and ‘platelet’ by two. The perfect score is zero, being the same node. This resembles that superclasses are usually more reliable, and have more in common than partly related, specialized subclasses.

## 3 Results

### 3.1 Database Sizes

An overview of currently contained annotations (as of 18.01.2024) in the database is shown in Table 2. It shows a comparison between the manually curated CellMarker2, and PuMA. Interactions refer to all co-occurrences between cells and genes, based on their respective dataset.

**Table 2:** Comparison of entries in CellMarker2, and PuMA (as of 18.01.2024). Interactions for cells and genes are corresponding entries which have at least one gene/celltype-relation. This excludes any entries without corresponding relation in the database. Marked* entries are filtered with at least three referencing articles (see filtering row for details).

| Entry | CellMarker2 | PuMA |
| --- | --- | --- |
| Filtering | #PMIDs $\geq 3$ | see sec. 2.2 |
| Distinct Cells |  |  |
| total | 449 | 1 009 |
| interactions | 449 | 628 |
| filtered* | 398 | 628 |
| Distinct Genes |  |  |
| total | 22 037 | 49 336 |
| interactions | 22 037 | 19 615 |
| filtered* | 7247 | 19 615 |
| Interactions |  |  |
| total | 96 074 | 195 288 |
| filtered* | 3 813 | 195 288 |
| Annotated Articles<sup>1</sup> | 6 322 | 9 847 891 |

<sup>1</sup> Relevant processed articles with useful annotations; total screening number may differ substantially.

It can be observed that a high number of annotations is excluded for CellMarker2 when partially filtering is applied. PuMA contains the highest number of distinct cells (increase by 1.6x) as well as distinct genes (factor 2.7x) in comparison to the filtered CellMarker2 database. Of note, the applied filtering of PuMA is performed before being deployed to the user. The total number of cells and genes annotated is even higher, and additional articles may influence a broad variety of yet to be included co-occurrences. The total number of found interactions is substantially larger than CellMarker2’s counterpart with a filtered version about 50x larger. The number of annotated articles is also increased manifold. This outlines its size advantage compared to CellMarker2.

#### Annotation performance

Figure 3 outlines the occurrence of genes in either database for both real datasets. For reference, 274 additional genes are found in PuMA, but not in CellMarker2 for the PBMC dataset. This corresponds to **≈** 75% (274/(274+90)) of previously unannotated genes by CellMarker2. For the TM dataset, our database extends CellMarker2 by **≈** 36%**Fehler! Textmarke nicht definiert**. (134/(134+240)) of previously unannotated genes.

**Figure 3.**
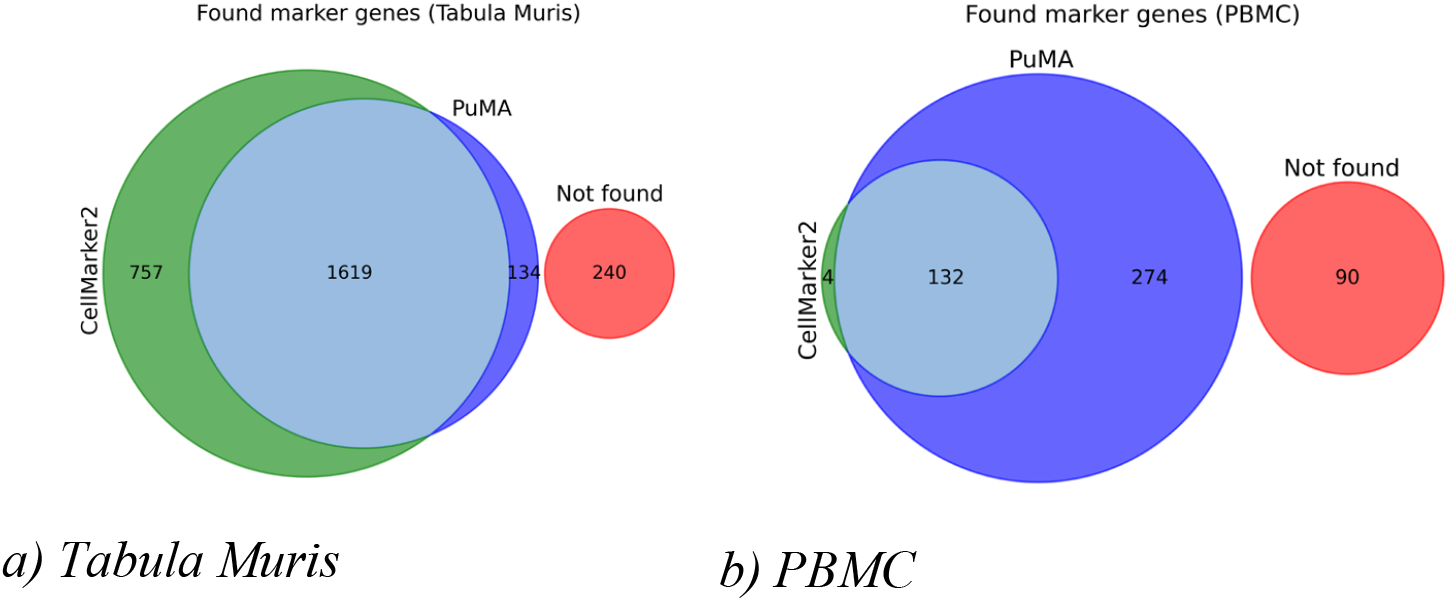
Matched number of top 50 DEGs in their respective databases for Tabula Muris (a) and PBMC (b). The red area (Not found) denotes marker genes not found in either database.

**Figure 4.**
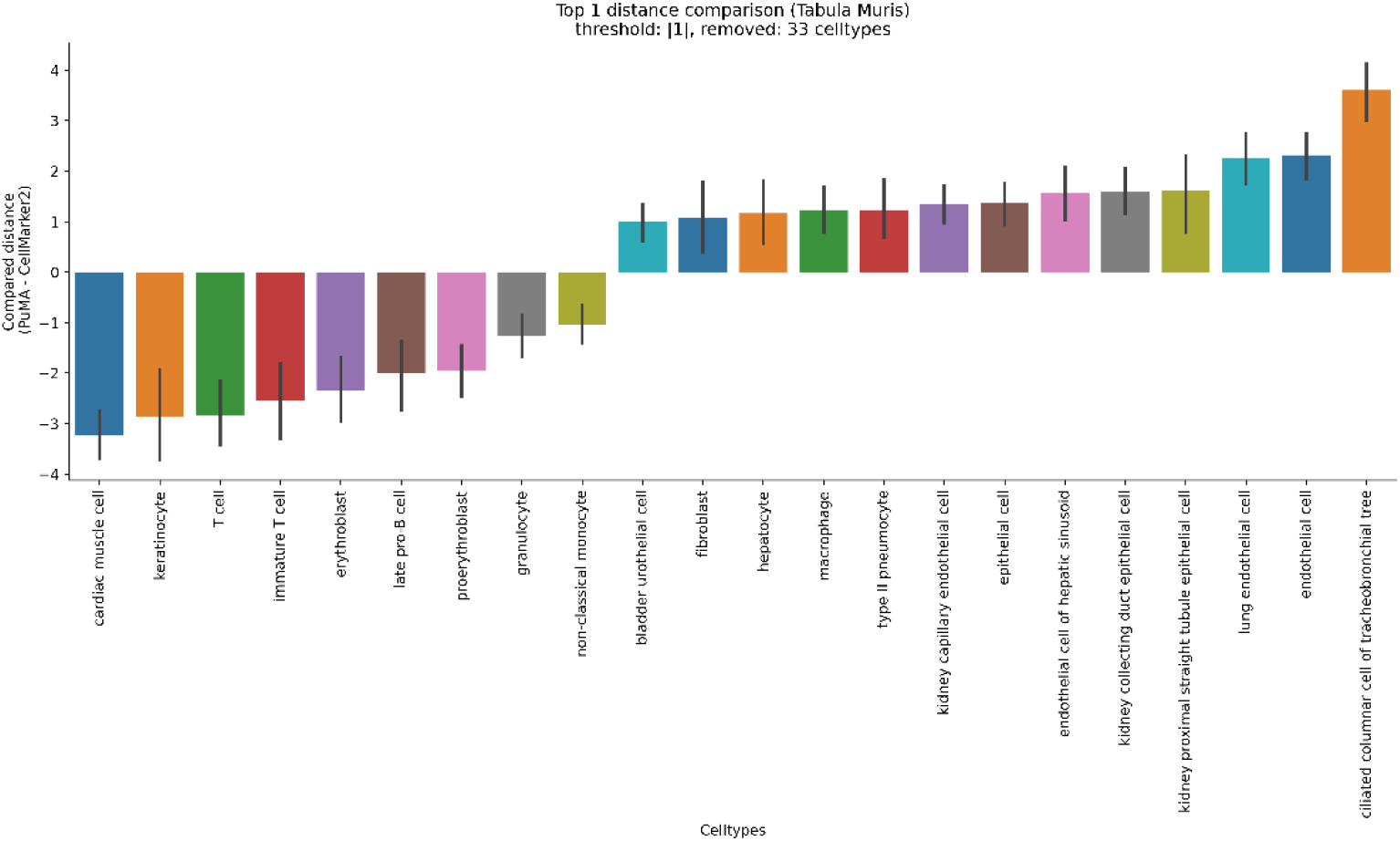
Per celltype compared distance performance for all genes (top-1 hits) of Tabula Muris. Lower values indicate better annotation performance of PuMA, while higher values suggest better performance of CellMarker2. A mean threshold of ±0.5 is applied for which these celltypes are removed from the plot.

#### Direct Comparison

Figure 4 and Figure 5 show the performance of PuMA vs CellMarker2 on the TM and PBMC dataset respectively. It stands out that PuMA achieves superior performance versus CellMarker2 for a number of celltypes, and is able to perform competitively to another set of celltypes in both datasets. As expected, CellMarker2 outperforms PuMA for a few celltypes, but mostly in the TM dataset. In general, we match or outperform CellMarker2 in top1 performance for more than 79.55% of matched genes in the PBMC dataset, and of about 62.26% in TM. The mean-distance, in a direct comparison between overlapping genes, decreases by **≈** −1.508 in PBMC and increases by **≈ +**0.0389 in TM respectively. In general, including all marker genes, PuMA has a median distance against CellMarker2 of −1.498 across all celltypes in the PBMC dataset, and +0.0015 in TM respectively. It performs better in the PBMC dataset than CellMarker2, and slightly worse in TM.

**Figure 5.**
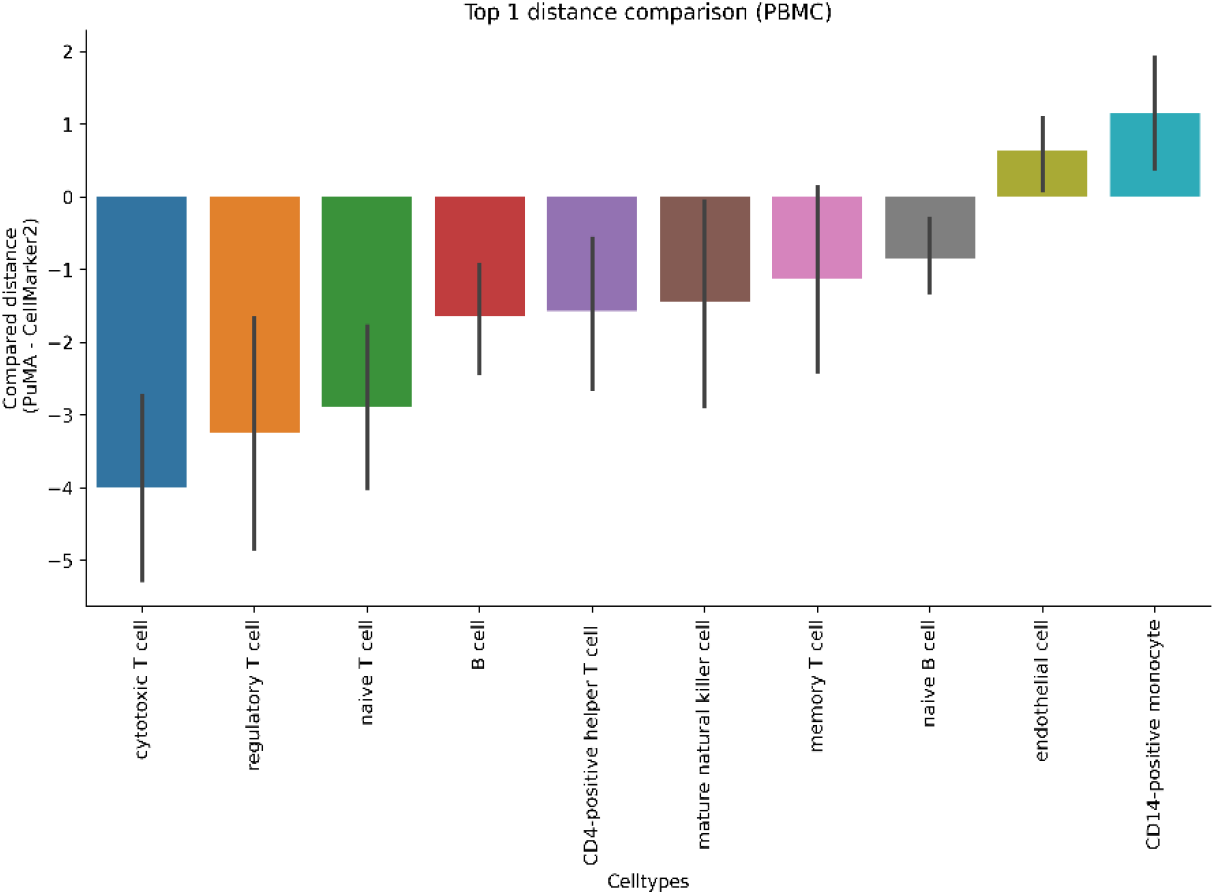
Per celltype compared distance performance for all genes (top-1 hits) of PBMC.

#### Combined Annotation Performance

Figure 6 and Figure 7 visualizes the combined annotation performance. The graphic plots CellMarker2 against the combined distance, which is the minimal distance of the top-1 hit in either database for all matched celltypes×genes:

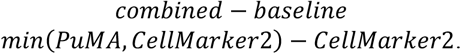

**Figure 6.**
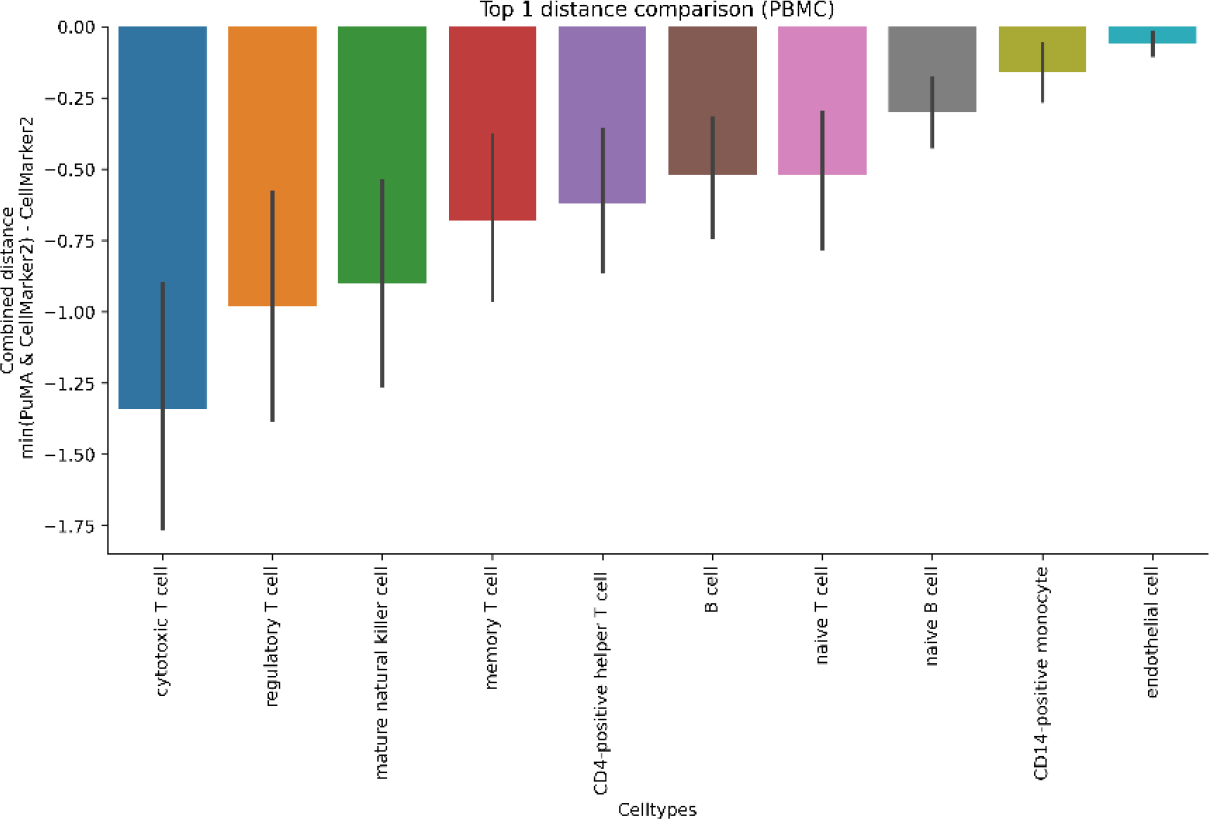
Annotation performance (top-1) for all genes for PBMC. The combined distance shows the combined average distance reduction for reference celltypes by using the minimum distance of PuMA or CellMarker2 respectively.

**Figure 4.**
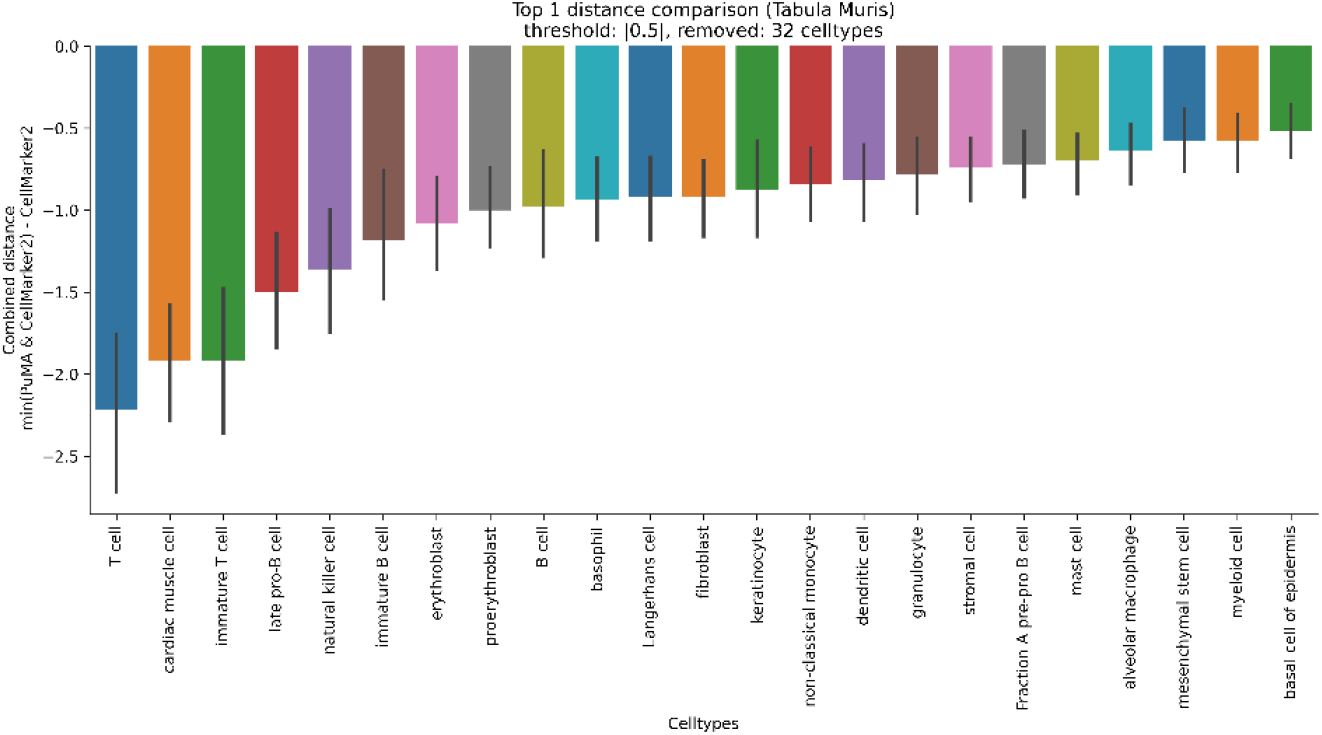
Annotation performance (top-1) for all genes for Tabula Muris. The combined distance shows the combined average distance reduction for reference celltypes by using the minimum distance of PuMA or CellMarker2 respectively. A mean threshold of ±0.5 is applied and these celltypes are removed from the plot.

It can be clearly distinguished that many annotations benefit from our database, underlining the importance of multiple data-sources. The combined mean distance reduction in comparison to plain CellMarker2, including all found genes in either database, can be reduced by 34.39% in PBMC, and up to 14.70% for TM.

## 4 Discussion

In this work, we presented PuMA, a novel, fully explainable database for gene/celltype-relations, which can automatically process PubMed articles, including recent literature. The web-based application provides a variety of tools, such as gene- or cell-search and rapid graph-based visualization. Especially cross-interactions between genes and different cell types can be interactively explored using PuMA. This may uncover previously unknown relations and empowers researchers to subsequently search for any related interactions. Additionally, we provide the user with all relevant articles to each result, which leads to an explainable tool with full traceability.

It can be argued that several well-established approaches identifying gene/celltype-relations already exist, some of them aiming at including recent publications. One example is CellMeSH (18), which has been created by processing MeSH-terms and Gene2Pubmed (8) as resources. However, CellMeSH relies on MEDLINE MeSH-cells and therefore can only cover a subset of PubMed content. Additionally, it does not include automatic updates, or the ability for computing these locally. Moreover, MeSH terms are broader and less distinct than its multiple counterparts in the Cell Ontology. While it is comparably large in the number of distinct cells and genes, it contains many unrelated annotations, which do not represent celltypes. These include cell names such as 54 variations of ‘chromosome’ and ten names containing ‘membrane’. The necessary additional filtering, and mapping to the open, accessible, and readily available mapped databases requires considerable curation work.

The compared datasets are commonly used datasets, and the evaluation is based on the reference celltypes. Also, the extracted marker genes are taken from Shunfu Mao et al., which do rely on the reference celltypes and provided biological data. We cannot fully ensure that biological truth is fully represented for all celltypes, and that the used clusters and its derived genes are completely distinct marker genes and there is no wrong clustering involved. Still, the evaluation is the same for PuMA and CellMarker2 and does not suffer from any advantage for either of these datasets.

While our database shows comparable performance for complete datasets, the quality varies by a gene-by-gene basis. However, if given enough marker genes, we can reach the competitive performance across datasets and meanwhile providing easy-to-use and additional tools to biologists. While CellMarker2 has a website available, it is hosted on an unencrypted server. Additionally, users are reliant on the server infrastructure to use its additional tools. While we provide docker locally deployable containers, not only for the web-interface, but also to update the dataset, this additional installation may hinder users to install it locally. We try to pursue detailed instructions to the user to provide them with necessary basic knowledge for a local installation.

Based on our results, we showed that PuMA is competitive against an extensive, up-to-date curated database, while including more marker genes and possible celltypes. In addition, we show that a combination of PuMA and CellMarker2 does bring incremental improvements, with a mean distance reduction regarding celltype similarity for both datasets at approximately 14.70% and 34.39% respectively, while being very competitive across both datasets and species. As PuMA can change with additional articles, we expect its performance to increase in the next few years, while new relevant articles are published on PubMed.

## 5 Conclusion

PuMA is a research tool for generating and visualizing gene and celltype relations by utilizing Large Language Models and PubMed articles. The explainability is facilitated through automated processing and linking to PubMed articles. PuMA performs competitively against a manual curated and up to date cell-marker database across two datasets and species. The combined approach of a high-quality manually curated database and PuMA extends matching genes and improves the overall annotation performance.

## Software and Code Availability

The software code of PuMA is open source, dockerized and available with installation instructions on: https://imigitlab.uni-muenster.de/published/PuMA.

